# Fluorescent tools for the standardized work in Gram-negative bacteria

**DOI:** 10.1101/2024.01.18.576257

**Authors:** Mario Delgadillo-Guevara, Manuel Halte, Marc Erhardt, Philipp F. Popp

## Abstract

Standardized and thoroughly characterized genetic tools are a prerequisite for studying cellular processes to ensure the reusability and consistency of experimental results. The discovery of fluorescent proteins (FPs) represents a milestone in the development of genetic reporters for monitoring transcription or protein localization *in vivo*. FPs have revolutionized our understanding of cellular dynamics by enabling the real-time visualization and tracking of biological processes. Despite these advancements, challenges remain in the appropriate use of FPs, specifically regarding their proper application, protein turnover dynamics, and the undesired disruption of cellular functions. Here, we systematically compared a comprehensive set of 16 FPs and assessed their performance *in vivo* by focusing on key parameters, such as signal over background ratios and protein stability rates, using the gram-negative model organism *Salmonella enterica* as a representative host. We evaluated four protein degradation tags in both plasmid- and genome-based systems and our findings highlight the necessity of introducing degradation tags to analyze time-sensitive cellular processes. We demonstrate that the gain of dynamics mediated by the addition of degradation tags impacts the cell-to-cell heterogeneity of plasmid-based but not genome-based reporters. Finally, we probe the applicability of FPs for protein localization studies in living cells using super-resolution microscopy. In summary, our study underscores the importance of careful FP selection and paves the way for the development of improved genetic reporters to enhance the reproducibility and reliability of fluorescence-based research in gram-negative bacteria and beyond.

## Introduction

Fluorescent proteins (FPs) have become indispensable tools for a broad spectrum of microbiological research and applications. The diverse repertoire of FPs enables *in vivo* monitoring of gene expression or protein-localization from bacterial populations to single cells. Transcriptional reporters usually comprise the fusion of a promoter of interest to the desired FP either *in cis* within the genome (in the native locus or ectopically) or *in trans* on a plasmid. When modifying the native locus of a gene of interest (GOI) to generate transcriptional fusions, FPs are inserted directly downstream of the coding sequence. Important considerations for any transcriptional reporter construct include the use of an appropriate ribosome binding site and spacer sequence to assure efficient transcription and translation^1,2^. Quantifying changes in the fluorescent signal of the reporter FP subsequently serves as a proxy for gene expression. Alternatively, an in-frame fusion between the GOI coding sequence with an FP on either C- or N-terminus of the protein of interest (POI), with or without a peptide linker, results in a translational reporter fusion enabling to study POI localization, dynamics, or turnover rates^3,4^.

In the past, comprehensive efforts have been made to optimize FPs to improve their brightness, photo-stability, and maturation time^5,6^. Perhaps the most commonly known green FP, GFP, originally from *Aequorea Victoria*, has been extensively engineered resulting in derivates with increased brightness and deviating excitation and emission characteristics^7^. For example, a mutational study on GFP yielded in an enhanced GFP (eGFP) variant with a 20-fold increase in fluorescence signal compared to the native precursor protein^7^. Additional engineering based on GFP also led to the discovery of reporter proteins with modified spectral characteristics, such as blue-shifted: mCerulean^8^ and mTurquoise2^9^ and yellow-shifted FPs: mVenus^10^, Ypet^11^. These achievements yielded in the development of a notable high fluorescent protein emitting in the green and yellow spectrum: mNeonGreen^12^. Additionally, the discovery of red FPs (RFP) extended the excitation range covered by reporter proteins within the spectrum of visible light^13^. Among these, mScarlet-I represents an optimized version of RFP, as it does not oligomerize and exhibits increased brightness, with reduced green maturation and excitation properties when compared to other RFPs such as mCherry or mRuby3^14^. The discovery of new FPs and their optimization has led to a repertoire of approximately 1,000 different FPs covering the spectrum of visible light^15^. In addition, to efforts driving the optimization of intensity and spectra specifications, fluorescent reporters have also been subject to engineering processes aimed to enhance their usability for specific purposes. A prominent example here is superfolder (sf)mTurquoise2ox, which was deliberately optimized to function outside of the bacterial cytosol and within the periplasmic space to serve as ideal Förster resonance energy transfer (FRET) partner^16^. Similarly, miniSOG was designed to bridge the diverse applications of correlated light and electron microscopy, with the positive side effect of resulting in one of the smallest FPs to date that require no additional co-factor^17^.

Compared to other frequently used reporter genes such as β-galactosidase (*lacZ*) or chloramphenicol acetyltransferase (CAT), FPs shine with main advantage to study cellular dynamic processes in real time without the necessity of endpoint measurements involving cell disruption^18,19^. For their functionality, FPs require no additional substrates except oxygen at the level of 0.1 ppm for chromophore maturation processes^20^. With their relatively small sizes of typically about 280-400 amino acids (AA), transcriptional or translational FP fusions are possible with minimal genetic modifications, which promotes their usability over larger reporters such as luminescence that is frequently based on the *luxABCDE* operon (of *Photorhabdus luminescens*)^21^. The flexible handling of FPs is also reflected by the versatile possibilities of readout options ranging from bacterial populations of colonies or in liquid cultures using plate-based or plate-reader devices, down to single-cells visualized by flow-cytometry or fluorescence microscopy.

Numerous studies in the past decades have proven FPs as primary choice for reporter genes, in particular to investigate transcription in time and space. Resolving phenotypic heterogenous traits of bacterial responses towards extrinsic cues or monitoring genetic cascade pathways and networks have previously been impossible to address adequately without the use of FPs^22–24^. In this light, a thorough examination and characterization of these reporters is indispensable, to assure appropriate reflection of even rapid occurring cellular processes.

In this study, we present an in-depth characterization of a comprehensive set of FPs covering the entire spectrum of visible light and provide examples of their *in vivo* applicability in bacterial cell biology using *Salmonella enterica* as a representative gram-negative host (Table 1, https://www.addgene.org/Marc_Erhardt/). *S. enterica* is a key model organism for studying pathogenic processes, motility behavior, metabolic pathways, and antibiotic responses and exhibits phenotypic heterogeneity in many of its pathogenic traits^25^. To enhance the usability of FPs for these areas, we first provided a thorough description of the optimal application of FPs, either plasmid-based or chromosomally encoded, followed by a detailed characterization of reporter dynamics. We investigated the use of degradation tags, both plasmid- and genome-based to determine their impact on reporter FP stability enabling investigation of temporal processes on shorter time scales. We corroborated our findings by assessing the constitutive expression levels of a ribosomal promoter throughout different growth phases, highlighting the importance of incorporating degradation tags for accurate tracking of expression dynamics at various growth phases. Finally, we assessed the use of FPs as translational fusions to investigate protein localization and dynamics. In summary, we envision that our findings will benefit the *Salmonella* community to conduct bulk or single-cell analyses, and empower the investigations of rapidly changing cellular processes.

**Table 1:** Collection of fluorescent tools provided in this study.

| FP* | Excitation<sup>1</sup> | Emission<sup>1</sup> | Addgene reference |
| --- | --- | --- | --- |
| mCerulean<sup>8</sup> | 452 | 479 | 213986 |
| sfmTurquoise2ox<sup>26</sup> | 455 | 478 | 213987 |
| sfGFP<sup>16</sup> | 492 | 512 | 213988 |
| mGFPmut2<sup>7</sup> | 489 | 509 | 213989 |
| eGFP<sup>7</sup> | 494 | 511 | 213990 |
| miniSOG<sup>17</sup> | 453 | 522 | 213991 |
| mVenusNB<sup>24</sup> | 517 | 530 | 213992 |
| Ypet<sup>11</sup> | 517 | 530 | 213993 |
| mNeonGreen<sup>12</sup> | 506 | 519 | 213994 |
| mRuby3<sup>27</sup> | 557 | 596 | 213995 |
| mScarlet-I<sup>14</sup> | 571 | 594 | 213996 |
| mCherry<sup>28</sup> | 585 | 611 | 213997 |
| mCherry2-L<sup>29</sup> | 583 | 613 | 213998 |
| mNeptune2<sup>30</sup> | 600 | 651 | 213999 |
| mNeptune2.5<sup>30</sup> | 597 | 639 | 214000 |

| Tagged FPs | Tag | Addgene reference |
| --- | --- | --- |
| mCerulean | LVA | 214003 |
|  | AAV | 214011 |
|  | ASV | 214012 |
| eGFP | LVA | 214001 |
|  | AAV | 214005 |
|  | ASV | 214006 |
| mNeonGreen | LVA | 214002 |
|  | AAV | 214010 |
|  | ASV | 214008 |
| mScarlet-I | LVA | 214004 |
|  | AAV | 214007 |
|  | ASV | 214009 |

\*Primary reference
<sup>1</sup>Excitation and emission optima experimentally obtained from spectra analyses (Figure S1).

## Results

### *In vivo* characterization of fluorescent proteins

To identify suitable candidates for the construction of transcriptional reporters, FPs covering the entire spectra of visible light were evaluated (Figure 1A). For this, 16 individual FPs were cloned into a plasmid downstream of the frequently used constitutive promoter (P_*rpsM*_) of the ribosomal protein S13 and introduced into *S. enterica*. Subsequently, the excitation and emission spectra of the FPs *in vivo* were recorded using a plate reader (Table S1 and Figure S1) and resembled previous *in vitro* measurements available in FPbase (https://www.fpbase.org/) (Table 1)^15^. Next, we performed endpoint measurements to compare the fluorescence intensities for all constructs over wild-type (WT) background levels normalized by optical density (Figure 1A). From this, we calculated the signal to background ratios and confirmed previous findings that mNeonGreen and mScarlet-I show a particularly high signal over background ratio, with a 1,000-fold increase in fluorescence over WT levels. Blue fluorescent proteins such as mCerulean depicted a 100-fold increase in signal to background, whereas sfmTurqouise2ox yielded only a 10-fold increase. In the green/yellow spectrum, mGFPmut2, Ypet, mVenusNB, and mNeonGreen show a 10-fold higher fluorescence intensity compared to other FPs within the same emission range (Figure 1B). Bacteria exhibit only weak autofluorescence at wavelengths above 500 nm (Figure 1A), which explains the relative high signal over WT background of FPs such as mCherry and mScarlet-I.

**Figure 1:**
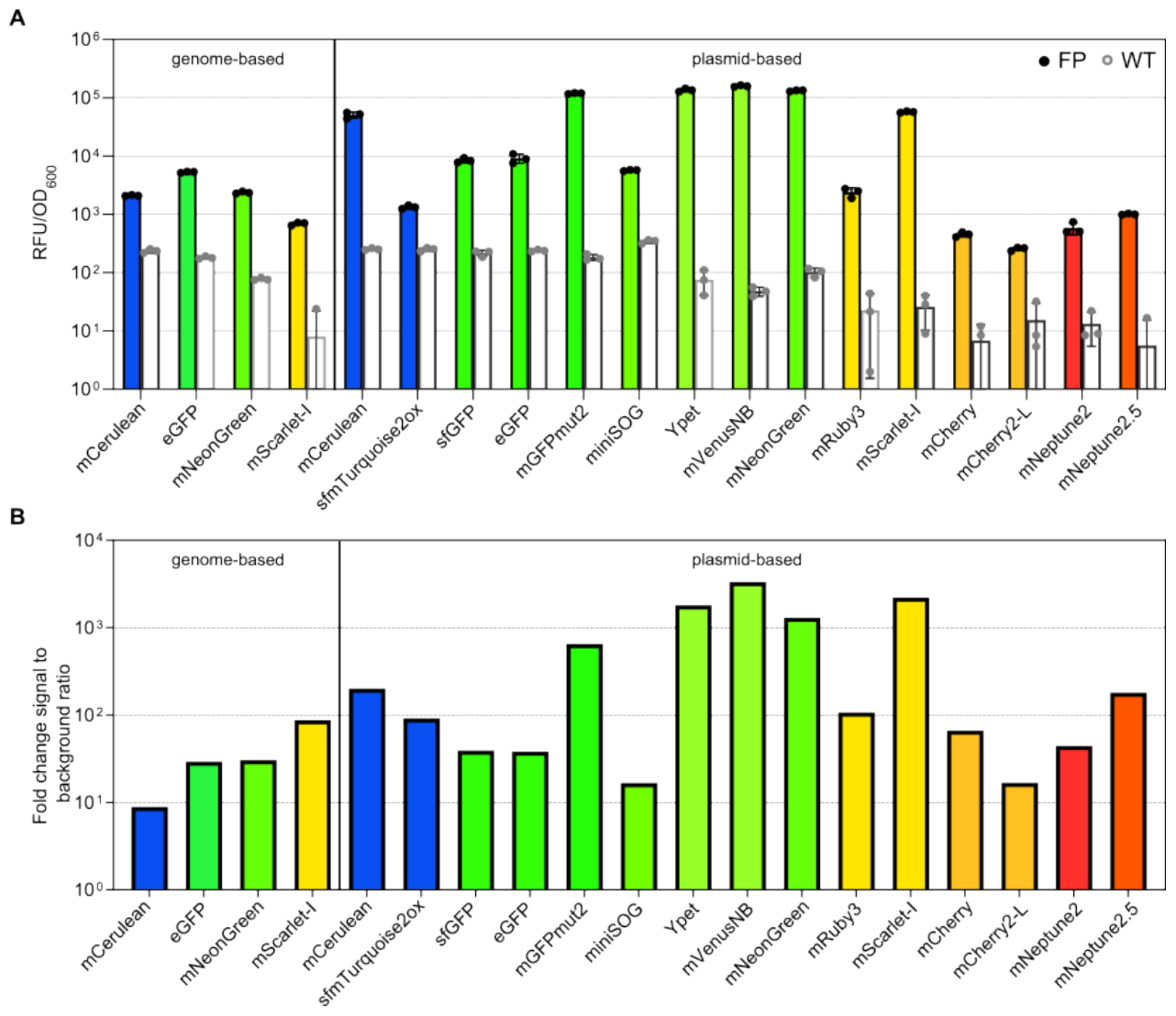
Endpoint measurements of the fluorescent proteins evaluated in this study, both genome- and plasmid-based. **A)** Relative fluorescent units normalized by optical density (OD_600_) are shown for cells constitutively (P_*rpsM*_) expressing the corresponding FP (colored bars), compared to the autofluorescence of *S. enterica* WT cells (white bars). The excitation and emission wavelengths for each FP are depicted in Table 1. Bar graphs represent mean values and standard deviations of at least three biological replicates. For evaluation of the spectra (Fig. S1) and endpoint measurements, cells were grown to exponential phase and fluorescence was directly measured in growth medium within a final volume of 200 μl in 96 well plates using a Synergy H1 plate reader. **B)** Fold-change of reporter signal normalized to WT background fluorescence depicted for each FP.

One major advantage of plasmid-based reporters is the ease of transforming a single construct into various backgrounds. However, plasmid-based reporters may skew the activity of a GOI because of enhanced gene dose due to copy number of the plasmid backbone^31^. Plasmid replication and reporter protein expression might also affect bacterial growth due the increased metabolic burden^32^. Lastly, plasmids may be non-homogenously distributed during exponential growth, when daughter cells receive different number of plasmid copies, rendering observations of phenotypic heterogeneity more difficult^31^. To overcome these limitations, native or ectopic transcriptional fusions are preferable despite a reduction in signal strength due to decreased gene doses. Accordingly, we next conducted further analyses using chromosomal constructs. For this, we chose representative green FPs (mNeonGreen and eGFP), one cyan (mCerulean), and one RFP (mScarlet-I). All reporters were constitutively expressed from the chromosome (replacing the non-essential *amyA* locus) and evaluated based on the excitation and emission optima determined using plasmid versions of the reporter constructs. First, we characterized the brightness of the chromosomally encoded versions via endpoint measurements in relation to the background fluorescence of the WT *S. enterica* (Figure 1A). Among the tested FPs, mScarlet-I showed the highest signal over background fluorescence with an 80-fold higher signal relative to wild type levels (Figure 1B). For both FPs in the green spectrum, mNeonGreen showed lower brightness than eGFP; however, the signal over the background fluorescence was slightly enhanced for mNeonGreen. Of the four tested FPs, mCerulean showed the lowest ratio compared to the WT, with only an 8-fold increase in signal over background.

After the general characterization of the FPs, we next focused on evaluating mCerulean, eGFP, mNeonGreen, and mScarelt-I in greater depth. For this purpose, we first manipulated the stability of the reporters to improve their applicability in monitoring rapidly changing cellular processes.

### SsrA-tag mediated degradation of fluorescent proteins to enhance reporter dynamics

Bacteria encode sophisticated protein control and quality systems to avoid synthesis of error-prone polypeptides. In *Escherichia coli* (*E. coli*) and *S. enterica*, the SsrA-tag system is utilized to release stalled ribosomes of transcripts lacking stop codons^33^. The *ssrA* RNA molecule, which functions as a transfer and messenger RNA (tmRNA), binds to stalled ribosomes. Upon binding of the tmRNA, a short open reading frame encoding the SsrA tag is translated and added to the nascent peptide. This leads to translation termination by the stop codon within *ssrA* and release of the stalled ribosome^34^. In turn, the peptide is now primed to be recognized by protease complexes including FtsH, Lon, Tsp, ClpAP, and most prominently ClpXP, resulting in the degradation of truncated and mistranslated proteins^34,35^. This system can be used to decrease the stability of the engineered proteins^36^. To explore the application of unstable reporter proteins and tune their temporal resolution capacity, plasmid-based constitutively expressed FPs were fused to different SsrA-tags with varying specificity to the degradation complexes (Figure 2A, B). This approach was previously described with degradation tags containing the amino acid sequences: AANDENYALVA (abbreviated as LVA), AANDENYAAAV (AAV), and AANDENYAASV (ASV) that displayed decreasing degradation efficiency in *E. coli*^37^. The variation in degradation efficiency depends on the affinity of the tag to host proteases. Subsequently, a high affinity to the protease mediates fast degradation, whereas a lower affinity results in slow degradation dynamics. Importantly, when comparing the degradation of identical SsrA-tagged GFP constructs between *E. coli* and *Pseudomonas putida*, the protein stability differed between the two species^37^. Thus, a species-specific characterization of such tags is indispensable. Therefore, we tested the degradation kinetics of representative FPs covering the entire spectrum of visible light using fusions of three variations of the SsrA tag in *S. enterica*. LVA, AAV, and ASV were individually fused to mCerulean, eGFP, mNeonGreen, and mScarlet-I, and the protein degradation kinetics were determined using a pulse-chase experiment. Translation was inhibited by the addition of chloramphenicol and spectinomycin to bacterial cultures grown to mid-exponential phase (Figure 2A, B). By comparing the fluorescence intensity of the tagged and non-tagged FPs at the time point of growth arrest (t=0), we observed that the presence of any SsrA tag increased the steady-state degradation of the FP, resulting in a reduction in the overall reporter signal strength (Figure 2A). LVA-tagged FPs decreased the signals of mCerulean, eGFP, and mScarlet-I by approximately 50%, whereas the signal of mNeonGreen was reduced by approximately 85% compared to the untagged FPs. The AAV and ASV tags decreased the reporter strength of mCerulean, eGFP, and mScarlet-I by approximately 40% and 10%, respectively. Notably, AAV-tagged mNeonGreen depicted a reduction of about 70% and ASV-tagged mNeonGreen of 40% compared to the untagged FP (Figure 2A). After the addition of antibiotics and inhibition of *de novo* protein synthesis, we next investigated the tag-dependent degradation kinetics of the FPs by monitoring changes in the fluorescence intensity over a period of 3 h. For this, we determined the time at which 25% (T_25_) or half of the signal (T_50_) was reached compared to non-tagged FPs. Consistent with steady-state degradation levels, we observed that LVA-tagged proteins showed the strongest and fastest degradation kinetics (Figure 2B and Table S2). The AAV and ASV tags resulted in degradation kinetics between non-tagged and LVA-tagged FPs. Strikingly, for mNeonGreen but not for the other tested FPs, we did not observe differences in the degradation kinetics between the AAV and ASV tags (Figure 2B). For mScarlet-I, we detected an increase in the fluorescence signal for all constructs after inhibition of translation. The most likely explanation for this observation is a delayed fluorophore maturation after protein synthesis. A similar yet less pronounced effect was observed for eGFP (Figure 2B), and the addition of any SsrA degradation tags reduced the overall signal strength for both FPs, demonstrating their general functionality in protein degradation.

**Figure 2:**
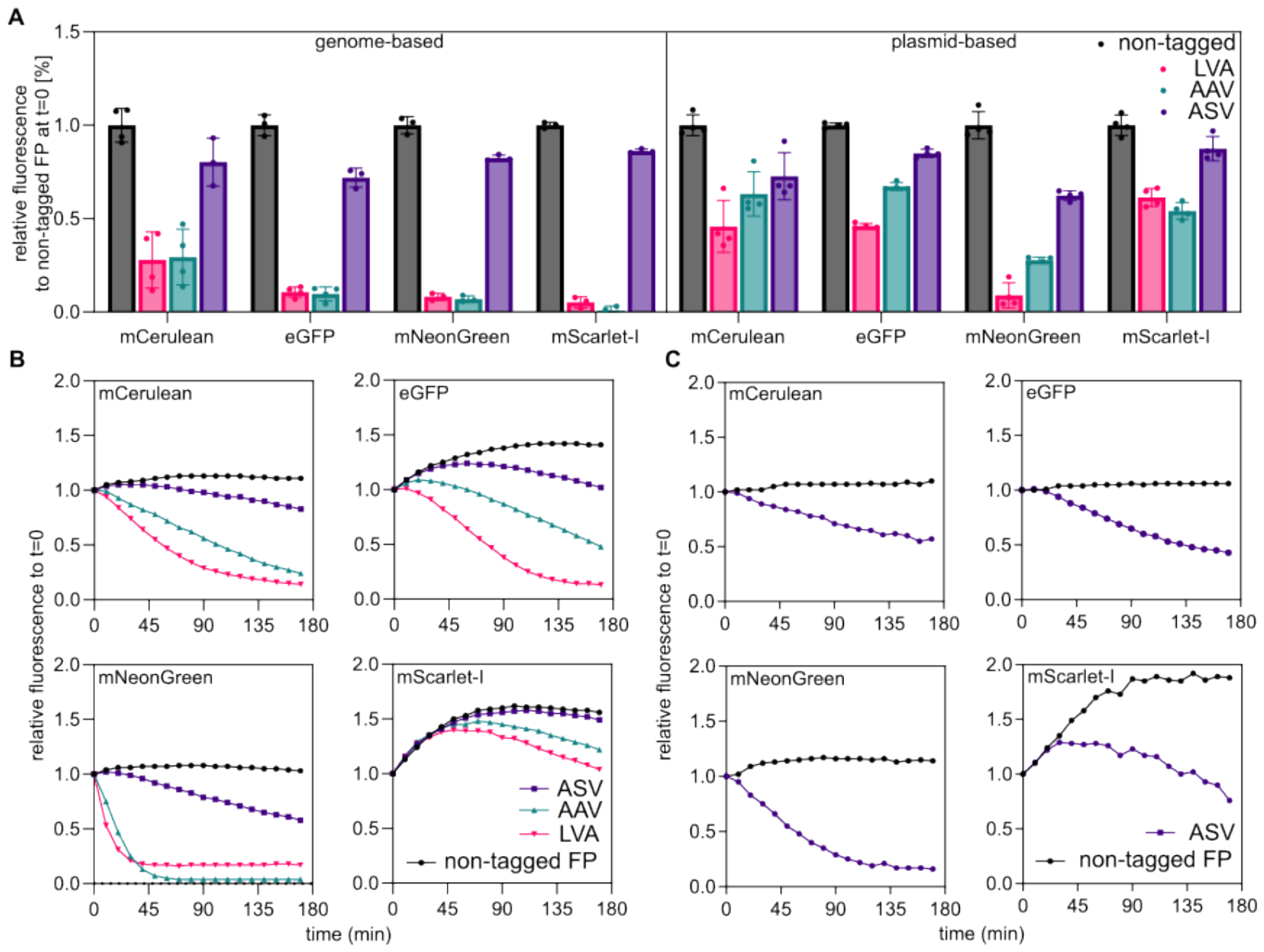
Evaluation of SsrA degradation tags that modulate the stability of FPs. **A)** Relative fluorescence intensity of tagged FPs compared with their non-tagged counterparts. The tags cause a mild to moderate drop of maximum FP intensity that correlate with their affinity to be degraded. **B+C)** Pulse-chase experiments to observe degradation speed over a time of 3 h after inhibition of *de novo* translation. Non-tagged FPs remain stable or show an increase in fluorescence intensity, due to maturation time of the FPs, the tagged FPs decrease in intensity at different speeds reflecting their degradation efficiency. For genome based tagged FPs, only fluorescence was detected for the ASV tag (see main text).

To the best of our knowledge, previous investigations concerning the degradation kinetics of SsrA tags have only been performed using plasmid-based reporters, which motivated us to characterize the degradation kinetics of genome-encoded FPs (Figure 2A, C). Surprisingly, for genome-encoded FPs, no fluorescence signal was detected for strains expressing FPs fused to tags mediating fast or intermediate degradation (LVA, AAV), independent of protein synthesis arrest, indicating efficient degradation before chromophore maturation (Figure S2). However, we observed a fluorescence signal in strains expressing proteins fused to the ASV tag, which mediated the slowest degradation within our set of degradation tags. When comparing the fluorescence of the non-tagged and tagged reporters at the time of antibiotic addition, mNeonGreen and mScarlet-I exhibited approximately 40% reduction in fluorescence intensity (Figure 2A). After inhibition of *de novo* protein translation, the levels of mNeonGreen-ASV decreased constantly, whereas for the non-tagged protein, the signal remained stable until the end of the experiment (Figure 2D). Similar to plasmid-based experiments, mNeonGreen displayed the fastest degradation with mNeonGreen-ASV, depicting a signal reduction to 50% within one hour after growth arrest. Three hours after translation inhibition, the fluorescent signal of mNeonGreen-ASV was reduced to 20% relative to non-tagged mNeonGreen. In comparison, the mScarlet-I signal initially increased after translation inhibition in the ASV-tagged and non-tagged FPs, as we previously observed for the plasmid-based version, and can be explained by a maturation delay of the already fully translated mScarlet-I prior to the addition of antibiotics. mScarlet-I displays a significantly longer maturation time than mNeonGreen, eGFP, or mCerulean^29^. Despite this maturation delay, the levels of mScarlet-I-ASV decreased by approximately 30% compared to the non-tagged reporter protein at the end of the experiment. Similar results were observed for the other two reporter proteins, mCerulean-ASV and eGFP-ASV, in which the fluorescence levels constantly decreased after translation inhibition (Figure 2A and C). After 3h of incubation, a decrease in fluorescence to approximately 40% and 20% was observed for mCerulean-ASV and eGFP-ASV, respectively. In summary, fusions of SsrA degradation tags to FPs decreased the stability of chromosomally encoded reporters, rendering these improved reporters more sensitive with an increased temporal resolution.

Since the proper function of the degradation tags is dependent on the activity of the host proteases, we next addressed how the deletion of ClpXP, a global protease complex, impacts the degradation kinetics of the reporter FPs^34,35^. For this, we separately transformed the plasmid-encoded eGFP reporter variants into a *S. enterica* ∆*clpXP* strain, performed a pulse-chase degradation experiment, and monitored reporter activity over time after translation arrest (Fig. 3A). As expected, the absence of ClpXP resulted in no degradation of ASV- and AAV-tagged eGFP compared to the non-tagged eGFP. Interestingly, we detected some degradation of the LVA-tagged construct (Fig 3A). These findings suggest that ASV- and AAV-tagged proteins are only recognized and degraded by ClpXP, whereas other proteases such as FtsH or Lon are capable of degrading LVA-tagged proteins in the absence of ClpXP^38,39^.

**Figure 3:**
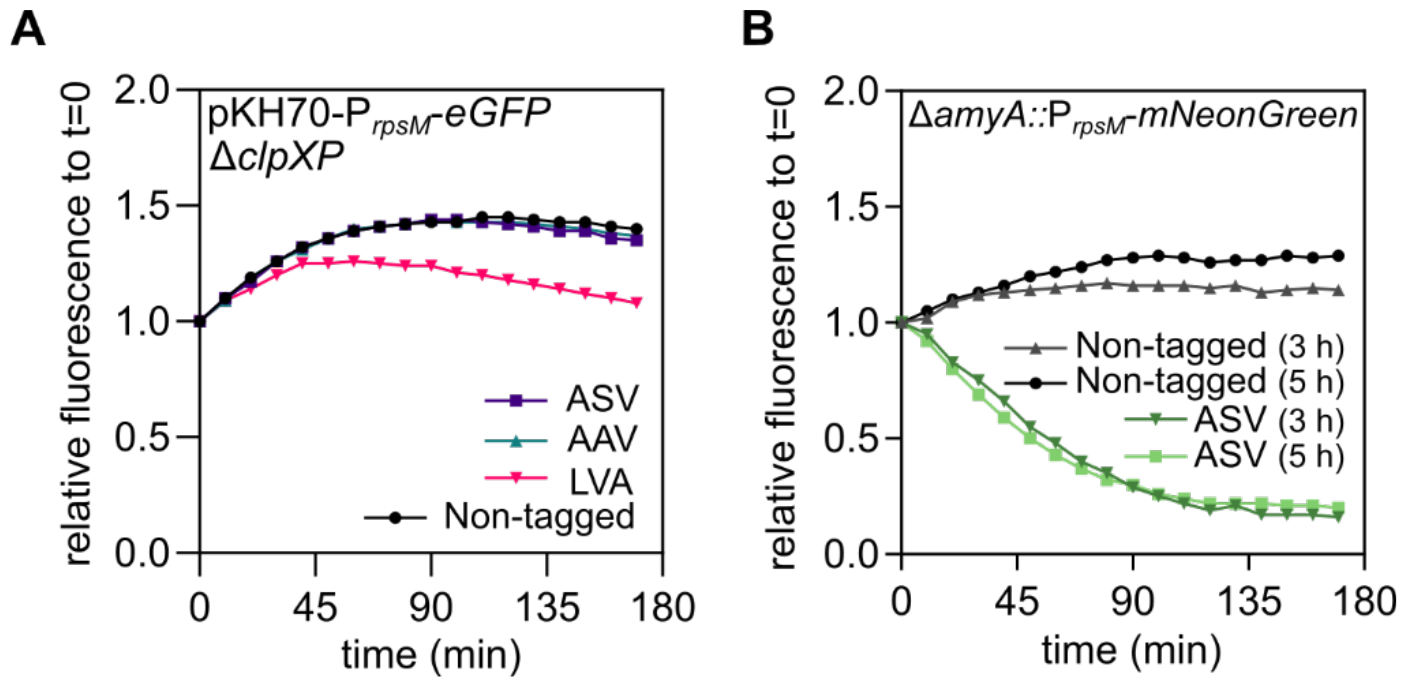
Effect of ClpXP on the SsrA-mediated degradation dynamics of tagged FPs. **A)** Pulse-chase experiment for three hours of plasmid-based eGFP with degradation tags or non-tagged in a ∆*clpXP S. enterica* background. **B)** Pulse-chase experiment of genome-based mNeonGreen expression either performed in exponential phase (3h) or stationary phase (5h).

In addition to the general importance of ClpXP for degradation of FP reporters, we next investigated how varying protease levels affect the activity of a SsrA tagged reporter construct at different growth phases. Based on the SalCom RNA sequencing database, we determined that *clpXP* transcription levels peak in mid-exponential phase and decrease substantially in early- and late-stationary phase^40^. Accordingly, we determined the degradation kinetics of a genome-based mNeonGreen reporter with and without ASV tag after 3h (exponential phase) and 5h (stationary phase) of growth (Fig. 3B). Interestingly, we did not observe any differences in the degradation kinetics, suggesting that at both tested timepoints sufficient levels of ClpXP were available to degrade the ASV tagged reporter (Fig. 3B).

### Tagged fluorescent proteins enhance temporal resolution of transcriptional fusions

In order to investigate the applicability of SsrA-tagged FP reporters for monitoring gene expression dynamics, we next probed the activity of the well-studied constitutive promoter P_*rpsM*_ that natively drives the expression of the ribosomal protein S13 throughout various growth phases^41,42^. Based on the above-mentioned global RNA sequencing database, we predicted strong and steady FP reporter levels expressed from P_*rpsM*_ ranging from early exponential phase into early stationary phase^40^. Subsequently, in late stationary phase, we predicted a notable drop of promoter activity due to the stringent response^43^.

Accordingly, we generated a transcriptional fusion of P_*rpsM*_ to mNeonGreen with or without ASV tag in the *amyA* locus of *S. enterica* and followed growth as well as reporter signal over a time course of eight hours (Fig. 4). Until late exponential phase (until about 3h after start of the experiment), we did not observe substantial differences in reporter activity between the tagged and untagged mNeonGreen variant. However, as soon as the bacteria entered stationary phase, the downregulaton of P_*rpsM*_ was only observed for the ASV-tagged mNeonGreen variant (Fig. 4). The non-tagged mNeonGreen construct displayed a constant signal over the entire time of the experiment, thereby masking the growth phase dependent transcriptional downregulation of P_*rpsM*_ (Fig. 4). This observation underscores the necessity of incorporating degradation tags to fully reveal promoter dynamics *in vivo*.

**Figure 4:**
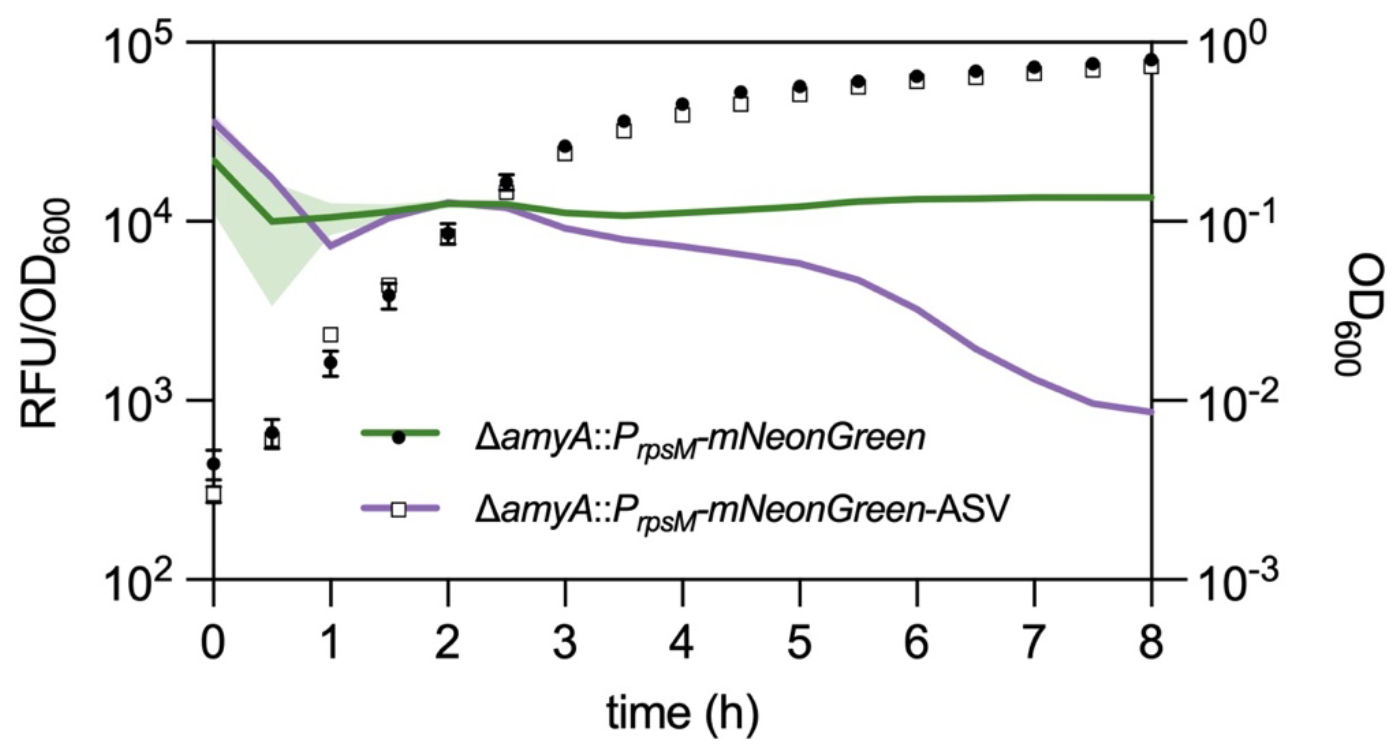
Transcriptional profile of tagged and non-tagged FP reporters. Growth and fluorescence intensity of strains harboring the ribosomal S13 promoter (P_*rpsM*_) fused to mNeonGreen or mNeonGreen-ASV were monitored in a plate reader over time. Growth and fluorescence measurements were performed every 30 min for eight hours. Means and standard deviation derive from at least three independent replicates and are represented by error bars for growth or shadowed space for fluorescence.

Next, we probed if the addition of degradation tags affects investigations of population heterogeneity. For this we analyzed the heterogeneity of genome-based and plasmid-based transcriptional fusions of P_*rpsM*_ to mNeonGreen with or without tags on the single cell level. As proxy for heterogeneity in gene expression, we determined the coefficient of variation (CoV) in fluorescence intensity of cultures grown to exponential phase using fluorescent microscopy (Figure 5). Surprisingly, a plasmid-based mNeonGreen lacking a degradation tag depicted a CoV of 0.31, which doubled for the LVA tagged version. Lower CoV values indicate less variation of measured fluorescence within a population set relative to the mean^44^. Satisfactorily, both the plasmid-based constructs harboring the ASV or AAV tag decreased the heterogeneous expression of mNeonGreen driven by P_*rpsM*_ compared to the non-tagged counterpart. This indicates that rapid degradation of a reporter fusion impacts cell-to-cell heterogeneity, and moderately strong degradation tags result in a more homogenous signal distribution, which should be considered while working with such constructs.

**Figure 5:**
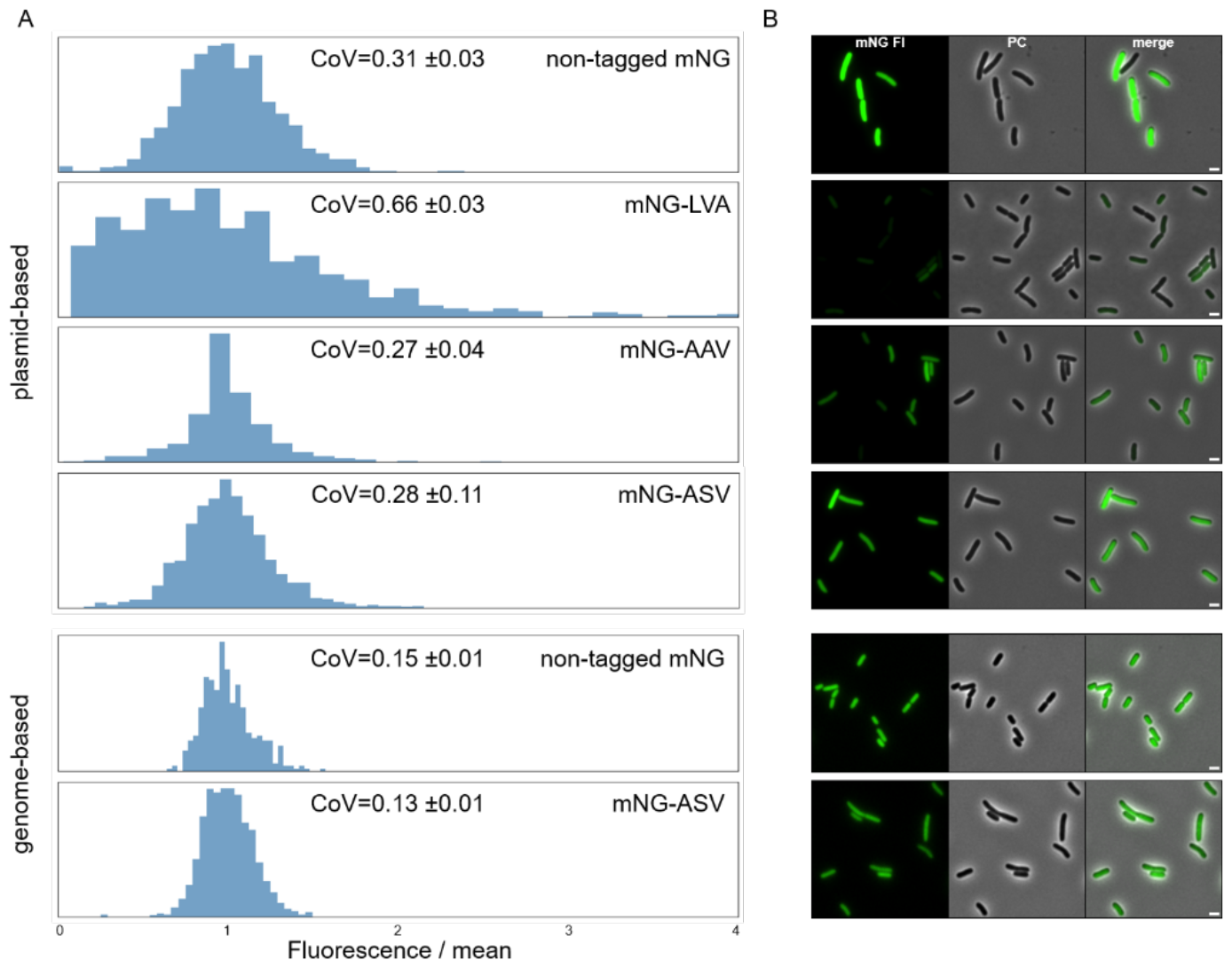
Determination of cell-to-cell variation using degradation tagged and non-tagged mNeonGreen transcriptional reporters. **A)** Coefficient of variation histograms of mNeonGreen driven by P_*rpsM*_ both on plasmid- as well as genome-based fused to one of the SsrA degradation tags or non-tagged. **B)** Exemplary fluorescence microcopy pictures of the constructs in A. Scale bar is 2 μm and the fluorescence signal intensities were adjusted separately for plasmid and genome-based images, respectively.

Although transcriptional reporters remain a major part of the application portfolio of FPs, translational FP fusions are also frequently used to investigate protein localization and dynamics. Therefore, we next set out to demonstrate the applicability of FPs as translational fusions in life cell studies, as well as fixed samples, using fluorescence and super-resolution microscopy.

### Fluorescence proteins to study protein localisation and dynamics

As a proof of concept, we fused representative FPs to a C-ring protein of the flagellar basal body in *S. enterica*. For this, we chose FliG, a protein part of the flagella type-III secretion system (fT3SS), which interacts with the stator units MotAB to drive rotation of the flagellum^45–47^. N-terminal, translational fusions were constructed at the native *fliG* locus (Figure 6A). To quantitatively evaluate the performance of FPs as proxies for FliG localization, we systematically analyzed the signal-to-noise ratio (SNR) in cells expressing each FP fusion using epifluorescence microscopy (Figure 6B). The SNR data indicated that while there was variability among the FPs, several candidates produced signals significantly above the background levels, which is suitable for localization imaging studies. For instance, sfGFP, mNeonGreen, and mScarlet-I exhibited higher SNR values compared to mCherry2-L or mNeptune2 fusions. However, fluorescence microscopy imaging revealed distinct localization patterns of the FPs corresponding to FliG complexes assembled in C-rings, and we observed that visualization of blue excitable FPs (sfmTurquoise2ox, mNeonGreen, sfGFP) was more easily detectable than orange/red excitable FPs and would therefore be more suitable for foci fluorescence observations (Figure 6C). Successful fusion and functional expression with the native POI locus require assessment of the functionality of the newly engineered protein. In the case of FliG, we assessed the impact of the fusions by confirming the presence of flagellar filaments and measuring overall motility. Although all fusions were flagellated, motility was affected differently depending on the fusion (Figure S3), indicating that the fusions did not prevent the secretion of flagellar subunits, but did affect interaction with stator units and therefore flagellar rotation^46^. We next utilitzed Stimulated Emission Depletion (STED) super-resolution microscopy to visualize mNeonGreen-FliG with a higher resolution (Figure 6D). We observed substantially more distinct FliG foce and reduced background using STED compared to confocal laser scanning microscopy (CLSM). Although FPs in general display lower photostability and are more prone to photobleaching than organic dyes, such as the HaloTag TMR ligand^48^, the increased resolution observed here demonstrates the potential of using selected FPs, such as mNeonGreen, for STED microscopy in bacteria. Collectively, these results underscore the versatile applicability of FPs also for the study of protein localization and functions in Gram-negative bacteria.

**Figure 6:**
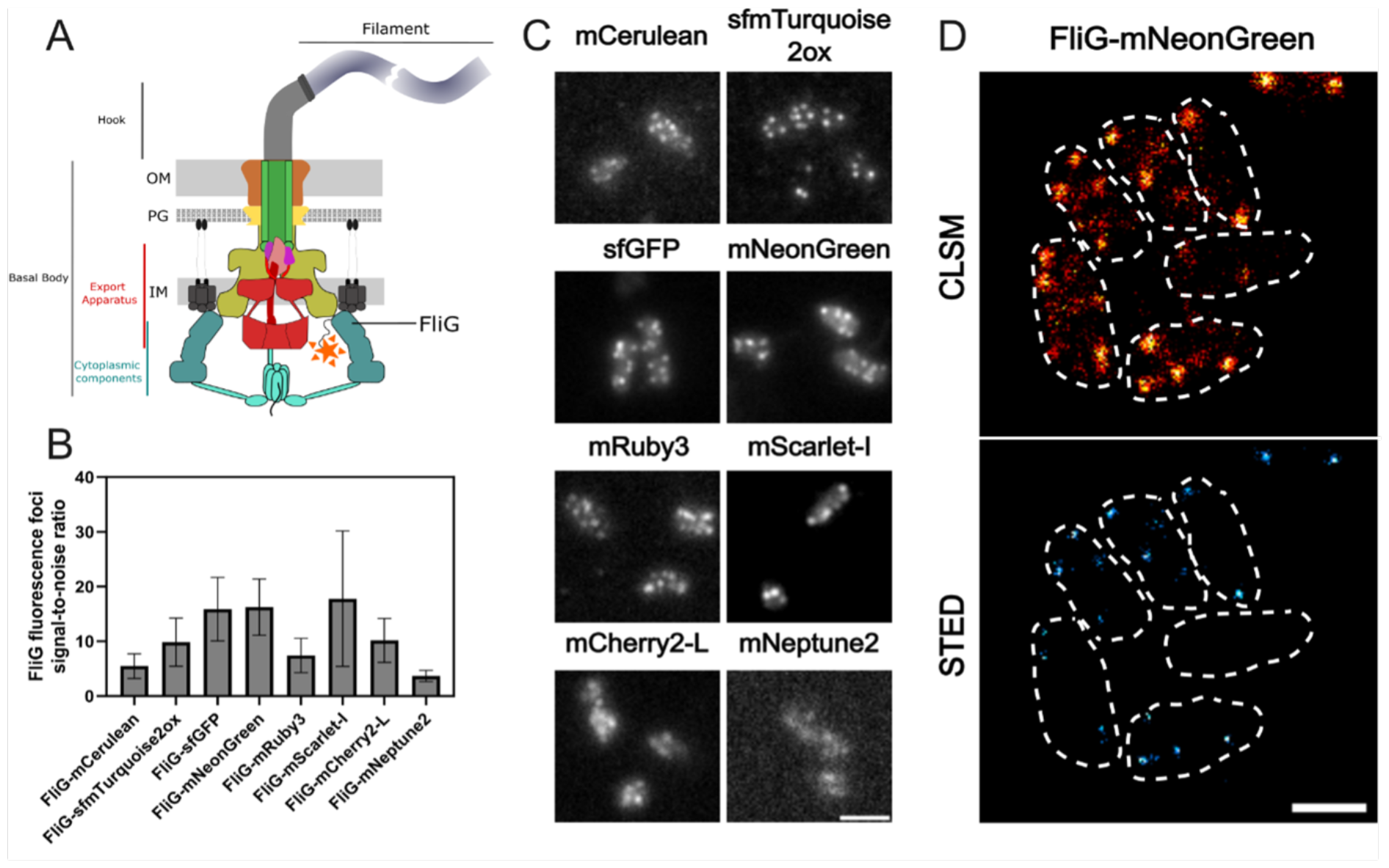
Visualizing cellular structures in bacteria using translational FP fusions **A)**Schematic representation of the bacterial flagellar motor highlighting the position of the FliG protein within the basal body structure. **B)** Histogram showing the signal-to-noise ratio of the FliG fluorescence foci of different fluorescent protein fusions, indicating the efficacy of each fluorophore for fluorescence focus observation. **C)** Representative epifluorescence microscopy images displaying the distribution of the FliG protein fused to various fluorescent proteins within the bacterial cells (scale bar = 2 μm). **D)** Representative confocal laser scanning microscopy (CLSM) images and super-resolution STED (Stimulated Emission Depletion) microscopy image of FliG-mNeonGreen fusion, showcasing the enhanced resolution of FliG protein localization using STED (scale bar = 1 μm).

## Conclusion

In this study, we provide an in-depth characterization of fluorescent proteins (FPs) as tools to study the dynamic and rapidly changing cellular processes, including protein localization, using *Salmonella enterica* as representative gram-negative host. Fluorescent proteins are pivotal state-of-the-art reporters in molecular microbiology, offering versatility in applications ranging from transcriptional to translational fusions with genes of interest. They enable real-time monitoring of transcription and protein localization within live cells. However, despite their widespread use, a comprehensive characterization of these tools, particularly in *S. enterica*, has remained largely unexplored. To address this gap, we evaluated a collection of plasmid-encoded FPs spanning the entire visible light spectrum (Table 1). Our findings on the emission and excitation properties of these FPs are consistent with those of previous studies, confirming their optimal functionality in the *S. enterica* model^15^. One of the challenges of using plasmid-encoded reporter systems is the variability of plasmid copies per cell, which can skew the dynamics of the reporter constructs. To mitigate this issue, we explored the use of chromosomally encoded reporters (Figure 1). These reporters, including mCerulean, eGFP, mNeonGreen, and mScarlet-I, offer broad excitation and emission spectra, and enable the combination of multiple FPs within a single bacterium. Chromosomally encoded reporters address several limitations of plasmid-based systems, such as cell-to-cell heterogeneity owing to plasmid copy number variations, plasmid instability, and the necessity of antibiotics for plasmid maintenance. However, the single-copy nature of chromosomal reporters often leads to reduced signal-to-noise ratios. This was evident in our study, where strains with a single copy of the reporter showed significantly lower fluorescence intensity than their plasmid-encoded counterparts. Next, we explored the use of reporters with reduced stability, enabling a more precise temporal resolution when monitoring transcriptional patterns. We utilized a system based on SsrA-tags, which are known to promote protease-dependent degradation of tagged proteins (Figure 2). Surprisingly, chromosomally encoded FPs tagged with fast-degrading SsrA sequences (AAV and LVA) were undetectable in pulse-chase experiments, underscoring the challenges of using fast-degrading tags for chromosomally encoded reporters. While plasmid-encoded systems with higher gene dosages allow for the maturation and detection of proteins targeted for rapid degradation, chromosomal reporters may undergo degradation before maturation. In contrast, a slow-degrading tag (ASV) allows a balance between protein turnover and detectability. We also investigated the influence of ClpXP on the degradation dynamics of tagged reporters. Deletion rendered AAV- and ASV-tagged eGFP indifferent to degradation, while the reporter fused to LVA was still susceptible to degradation. Satisfactorily, stationary grown cells that decrease *clpXP* expression were also able to substantially decrease reporter signal according to the degradation tags affinity. Investigating the expression patterns of a ribosomal promoter using a non-tagged FP compared to an ASV-tagged reporter further supported the necessity of adding the degradation tag. Only when increasing the turnover rate mediated by the ASV tag could the comprehensive dynamic profile be obtained across the growth phases of P_*rpsM*_ transcription (Figure 4). Complementary to the degradation dynamics, we also explored the impact of adding a tag to a reporter on the cell-to-cell heterogeneity of the reporter signal (Figure 5). By determining the coefficient of variance, we compared the reporter noise of exponentially grown cells expressing mNeonGreen at the single-cell level. We noted that plasmid-based LVA-tagged FPs skew the heterogeneity of promoter activity, whereas ASV- or AAV-tagged reporters result in a more homogenous distribution over non-tagged FPs. Finally, we provide insights into the application of FPs as translational fusions with a POI. We demonstrated that the tested constructs are physiologically functional and highlighted aspects of SNR using epifluorescence microscopy. Although FPs tend to bleach rapidly using super-resolution microscopy techniques, such as confocal or STED, we demonstrate that mNeonGreen is suitable for such applications.

In summary, we provided a comprehensive description of FPs that can be applied to gram-negative bacteria. We explored plasmid- and genome-based variants of these reporters and stated the impact of adding degradation tags by highlighting their drawbacks and advantages. Reporter dynamics can be increased by adding an LVA tag to the reporter on plasmid-based constructs, but the increased cell-to-cell heterogeneity in single-cell assays needs to be considered as a potential drawback. We recommend using an ASV tag, both for plasmid and chromosomal transcriptional fusions. Here, dynamic turnover rates were obtained, reflecting better the temporal dynamics of transcription while maintaining a homogenous signal within a bacterial population at the single-cell level. For translational fusions, mNeonGreen proved to be suitable for super resolution microscopy approaches. We hope that the descriptive and engineering experiences provided in this study will benefit future studies that rely on the use of FPs as reporters of choice in Gram-negative bacteria and beyond.

## Declarations

### Ethics approval and consent to participate

Not applicable.

### Consent for publication

All authors read the manuscript and provide consent for publishing.

### Availability of data and materials

Collection of tools described in this study are made available via Addgene: https://www.addgene.org/Marc_Erhardt/

### Competing interests

The Authors declare no competing interests.

### Funding

This work was supported by the European Research Council (ERC) under the European Union’s Horizon 2020 research and innovation program (agreement no. 864971) to M.E. MDG was supported by a Grand Challenge Initiative Global Health grant of the Berlin University Alliance (no. 113_MC_GH_MEL-BER_Erhardt_HU).

### Authors’ contributions

MDG and PFP conceptualized the study. MDG, MH and PFP performed experiments. All authors interpreted the data. MDG, MH and PFP wrote the first draft of the manuscript, which was then edited by ME and PFP. All authors contributed to the revision of the manuscript.

## Acknowledgements

We thank all current and past lab members of the Department of Molecular Microbiology at the Humboldt-Universität zu Berlin for fruitful discussions and feedback on this study.

## Methods

### Bacterial growth conditions

Strains used in this study derived from *Salmonella enterica* serovar Typhimurium strain LT2 and are listed in Table S3. Bacteria were grown at 37°C under constant shaking (180 rpm) in lysogeny broth containing 1% tryptone, 0.5% yeast extract and 5% (*w*/*v*) sodium chloride. *Salmonella* strains harboring reporter plasmids were selected on ampicillin (100 μg ml^-1^). For maintenance of λ-Red genes encoding plasmid (pWRG730), bacteria were selected on chloramphenicol (10 μg ml^-1^) and grown at 30 °C.

### Chromosomal modifications

Chromosomal mutations were obtained using λ-Red-mediated recombineering and a *Kan*-SceI cassette as selection marker. Site-specific chromosomal insertion of the *Kan*-SceI cassette and subsequently the replacement with the desired chromosomal mutation was achieved by inducing the expression of the λ-Red phage genes (gam, exo, beta) by heat-shock from the temperature-sensitive plasmid pWRG730^49^. Both, the *Kan*-SceI cassette and the replacement fragments were amplified by PCR with flanking homologous regions of 40 bp to the target region. The desired chromosomal modification was obtained by counter-selection on LB plates containing anhydrotetracycline to induce expression of the meganuclease I-SceI. Successful recombination was verified by PCR, followed by sequencing. The primers used for chromosomal modifications are listed in Table S4.

### Plasmid cloning via restriction enzymes

Plasmids encoding for tagged and non-tagged FPs were cloned via restriction enzyme cloning. In the example of pKH70-P_*rpsM*_-*eGFP* cloning, the *eGFP* gene was amplified by PCR with oligonucleotides to contain an overhang with the restriction site for NdeI and XbaI. The PCR product and the vector were enzyme-digested using NdeI and XbaI (New England Biolabs) for 2 hours. pEM10687 (pKH70-P_*rpsM*_-RBS-*mNeptune2*) was chosen for vector isolation, as cells containing the undigested plasmid can be easily discerned by the purple coloration of the colony. To avoid re-ligation between the digested vector and the excised mNeptune2, the product of the digestion reaction was dephosphorylated by the Quick CIP (New England Biolabs). For ligation of the vector and the PCR product, a ligation reaction was conducted using the T4 DNA ligase (New England Biolabs) with a vector-to-insert ratio of 1:3. The ligated product was transformed into *S. enterica* and transformants were selected on ampicillin-supplemented solid LB (100 μg ml^-1^). The primers used for cloning are listed in Table S4.

### Spectral characterization of fluorescence proteins

The spectral properties of plasmid-encoded FPs were characterized *in vivo* in *S. enterica*. Overnight cultures of bacteria containing plasmids encoding for constitutively expressed FPs were diluted 1:100 in fresh LB supplemented with ampicillin (100 μl ml^-1^) and grown for 2.5 h. After incubation, bacteria from a 1.8 ml sample were harvested by centrifugation (11,000 × g, 2 min, room temperature) and resuspended in 1× E-salts buffer. Bacteria were pelleted once more and resuspended in 750 μl 1× E-salts buffer. For determination of emission and excitation properties, 150 μl of the resuspended cells were distributed in a black 96-well plate with clear bottom (Grainer Bio). Emission and excitation spectra were measured using the setting shown in Table S1. At the same time, the OD_600_ was determined for the correction of fluorescence intensity to bacterial density.

Additionally, to compare the signal-to-background ratio between different chromosomally encoded fluorescent proteins, endpoint measurements at the optimal excitation wavelength for each protein were performed. For the determination of the cell’s background fluorescence, the fluorescence intensity and OD600 were measured in the wild-type S. enterica. Similar to the characterization of the emission and excitation spectrum, cells were grown for 2.5 h, washed with 1× E-salts buffer, and 150 μl of the cell suspension was distributed in a black 96-well plate. Emission and excitation setting are shown in Table S1. Measurement of emission was performed using the Synergy H1 microplate reader. Measurements were conducted in at least two technical replicates of at least three biological replicates.

### Fluorescence protein stability

To assess the dynamics in protein degradation of SsrA-tagged FPs, the dynamics in fluorescence of bulk cultures were determined using the Synergy H1 microplate reader. For this, 200 μl LB in a 96-well plate was inoculated with single colonies of the desired strains. On the following day, the cultures were diluted 1:100 in fresh LB and 100 μl were distributed in technical duplicates in a black 96-well plate with transparent bottom. Initially, bacteria were grown in the microplate reader at 37 °C under constant shaking. During this time, fluorescence intensity and OD_600_ were measured every 30 min for 3 h. To inhibit the synthesis of new FPs after initial growth, chloramphenicol and spectinomycin were added to the culture to a final concentration of 10 μg ml^-1^ and 50 μg ml^-1^, respectively. Following the addition of the translation-inhibiting antibiotics, fluorescence intensity and OD_600_ were measured every 10 min for 3.2h. Emission and excitation parameters were set for the endpoint measurements as shown in Table S1.

### Monitoring gene expression in bulk culture

Expression of P_*rpsM*_ was characterized in bulk culture by monitoring changes in the fluorescence intensity during cell growth in a microplate reader (Synergy H1). Day cultures of at least three biological replicates were prepared by inoculating 200 μl LB with a single colony of the desired strains in a 96-well microplate. Cells were grown for 8 h at 37 °C under aeration. Subsequently, cell suspensions were diluted 1:100 in a two-step 1:10 serial dilution in fresh LB. 150 μl of the diluted culture were distributed in a 96-well black microplate in technical duplicates. OD_600_ as well as fluorescence intensity changes were recorded every 30 min for eight hours. Fluorescence intensity was tracked for mNeonGreen using the setting shown in Table S1. To reduce sample evaporation, wells surrounding the bacterial dilutions were filled with water.

### Fluorescence microscopy to characterize gene expression heterogeneity

*Salmonella* strains were grown overnight at 37°C and 180 rpm in LB with the addition of 100 μg ml^-1^ of ampicillin if applicable. Subsequently, a day culture was inoculated 1:100 in five ml LB in the presence of the antibiotic for plasmid-based constructs and incubated as the overnight culture for 2.5 hours (OD_600_ ranged between 0.6-0.7). Two μl of cells were then individually transferred to a 1.2 % agarose pad (Invitrogen Ultrapure Agarose in MQ) and imaged. Expression of P_*rpsM*_ was characterized using a Nikon Eclipse Ti2 inverted microscope and a 60x CFI Plan Apochromat DM 60x Lambda oil Ph3/1.40 objective. For strains harboring plasmid-based mNeonGreen constructs LED 488 nm was set to 10% intensity and exposure time to 10 ms. For genome-based constructs the latter was set to 100 ms. Image analysis was performed using Fiji^50^ equipped with the plugin MicrobeJ^51^. Data derives from three biological replicates and final graphs were prepared using customized Python scripts run in Jupyter Notebook. Calculation of the coefficient of variance (CoV) was determined for each replicate individually.

### Motility test of FliG translational fusion

*Salmonella* strains were grown overnight at 37°C and 180 rpm in LB. 2 μl of overnight cultures were inoculated into soft-agar swim plates containing 0.3% agar followed by incubation at 37°C for approximately 4-5 h.

### Fluorescence microscopy to observe translational fusion and flagellation

*Salmonella* strains were grown overnight at 37°C and 180 rpm in LB. Subsequently, a day culture was inoculated 1:100 in five ml LB and incubated as the overnight culture for 2.5 hours. Cells were then loaded to a custom-made flow cell incubated in 0.1% Poly-L-Lysin^52^. The slide and cover slip were assembled in the presence of a double layered parafilm as a spacer. Cells were fixed with 4% paraformaldehyde for 10 min at room temperature, followed by washing with PBS and incubation for 10 min with 10% BSA as blocking agent. The flagellar filaments were immunostained by incubating with a α-FliC primary antibodies (Difco) diluted 1:1,000 in 2% BSA solution for 1 hour. Samples were subsequently washed with PBS and incubated with 10% BSA, followed by incubation with a secondary α-rabbit antibody conjugated to Alexa Fluor 488 or Alexa Fluor 555 (Invitrogen) diluted 1:1,000 in PBS for 30 min. Samples were washed with PBS and were mounted in Fluoroshield (Sigma-Aldrich). Slides were imaged using a Nikon Eclipse Ti2 inverted microscope and a 60x CFI Plan Apochromat DM 60x Lambda oil Ph3/1.40 objective, with a Z-stack comprising 5 sections every 0.4 μm.

### STED microscopy of FliG translational fusion

*Salmonella* strains were grown overnight at 37°C and 180 rpm in LB. Day culture was inoculated 1:100 in five ml LB and incubated as the overnight culture for 3 hours. 500 μL of the cultures were then washed twice with PBS (2,500 *x g*, 2 min) and 5 μL were applied to a 1% agarose pad in PBS. Samples were imaged with The Abberior STED Facility line microscope (Abberior Instruments GmbH) equipped with an UPLSAPO 100 × oil immersion objective lens (NA 1.4) and a Motorized xy stage (Olympus IX3-SSU). Excitation was achieved with pulsed diode lasers PDL-T 488 and were depleted using a 595 nm laser (PRFL-P-10-595-B1R). Pinhole size was set to 1AU. Gated detection was applied and the Abberior Rescue module were used. Images were acquired using the Abberior software Imspector.

### Image analysis

Image analysis was performed using Fiji^50^ equipped with the plugin MicrobeJ^51^. Data derives from three biological replicates and final graphs were prepared using customized Python scripts run in Jupyter Notebook or GraphPad Prism version 10.

## Notes

### Competing Interest Statement

The authors have declared no competing interest.

