## Supplement files for "Fluorescent tools for the standardized work in Gram-negative bacteria"

### Supplemental information

**Table S1: Spectra ranges for endpoint measurement determination of FPs used in this study**

| Settings for measurements | | | | | | |
|---|---|---|---|---|---|---|
| | Endpoint | | Excitation spectrum | | Emission spectrum | |
| FP | Excitation | Set emission | Set Emission | Range | Set Excitation | Range |
| mCerulean | 452 | 482 | 515 | 300-485 | 408 | 438-700 |
| sfmTurquoise2ox | 455 | 485 | 515 | 300-485 | 408 | 438-700 |
| sfGFP | 492 | 522 | 550 | 300-520 | 460 | 490-700 |
| mGFPmut2 | 489 | 519 | 550 | 300-520 | 460 | 490-700 |
| eGFP | 494 | 524 | 550 | 300-520 | 460 | 490-700 |
| miniSOG | 453 | 522 | 541 | 300-511 | 422 | 452-700 |
| mVenusNB | 517 | 547 | 570 | 300-527 | 492 | 522-700 |
| Ypet | 517 | 547 | 570 | 300-527 | 492 | 522-700 |
| mNeonGreen | 506 | 536 | 557 | 300-527 | 481 | 511-700 |
| mRuby3 | 557 | 596 | 632 | 300-602 | 533 | 563-700 |
| mScarlet | 571 | 601 | 632 | 300-602 | 533 | 563-700 |
| mCherry | 585 | 615 | 650 | 300-620 | 564 | 594-700 |
| mCherry2-L | 583 | 613 | 650 | 300-620 | 564 | 594-700 |
| mNeptune2 | 600 | 651 | 691 | 300-661 | 574 | 604-700 |
| mNeptune2.5 | 597 | 639 | 691 | 300-661 | 574 | 604-700 |

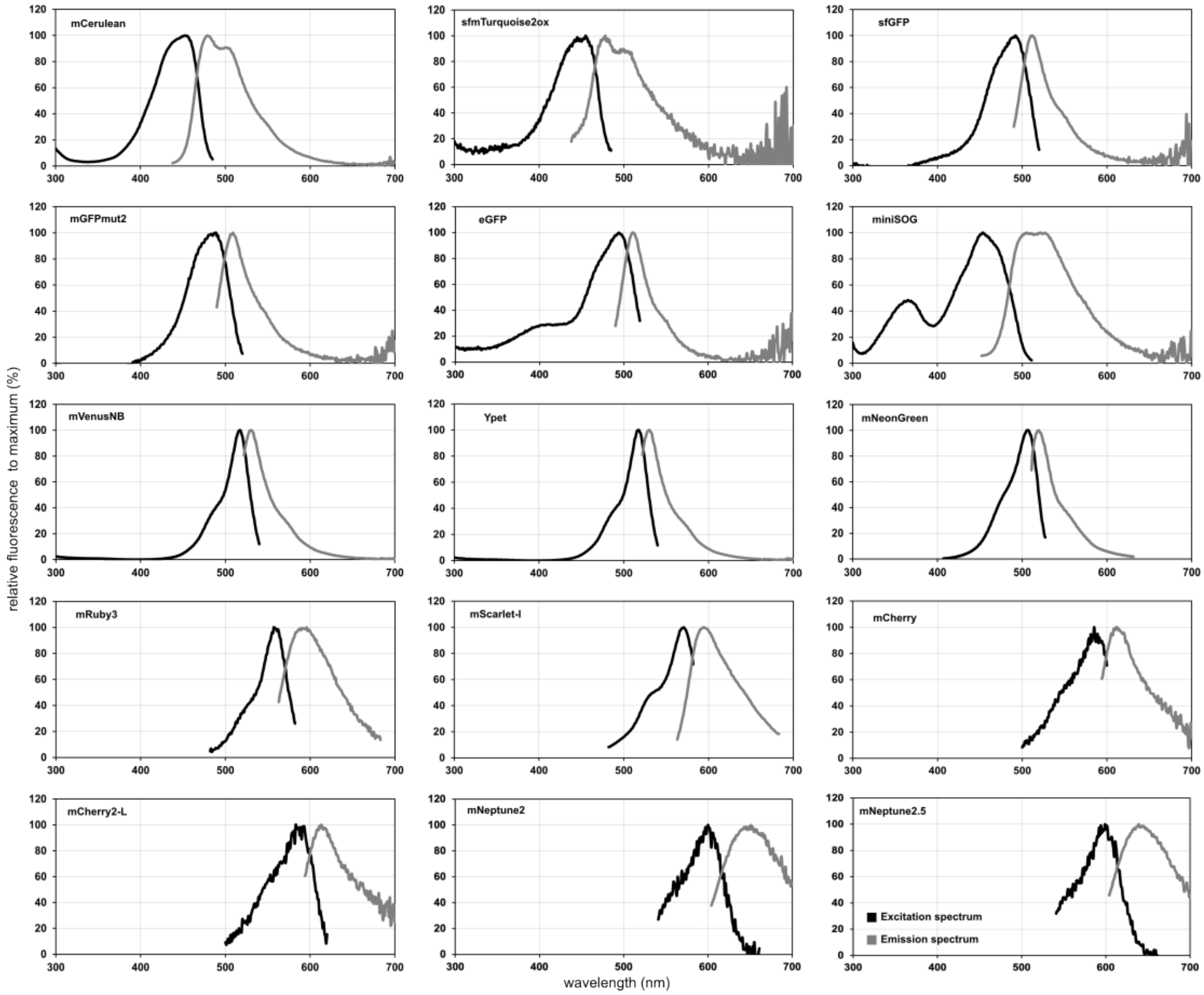


**Figure S1 Excitation and Emission spectra of FPs used in this study.**
Strains grown into exponential growth phase, harboring plasmids encoding respective FP driven by $P_{rpsM}$ were probed for their excitation and emission based on the nm ranges stated in table S1 using a Synergy H1 plate reader.

### Table S2 Half-life determination of FPs SsrA degradation kinetics

| FP | Tag | $T_{25}$ (min) | $T_{50}$ (min) |
|---|---|---|---|
| mCerulean | LVA | 29.4 | 55.2 |
| | AAV | 53.7 | 102.1 |
| | ASV | n.d | n.d. |
| | ASV (chromosomal) | 79.1 | 206.7 |
| eGFP | LVA | 49.0 | 74.2 |
| | AAV | 113.4 | 170.2 |
| | ASV | n.d | n.d. |
| | ASV (chromosomal) | 67.1 | 134.3 |
| mNeonGreen | LVA | 5.9 | 11.0 |
| | AAV | 10.5 | 18.0 |
| | ASV | 105.2 | n.d. |
| | ASV (chromosomal) | 30.6 | 55.6 |
| mScarlet | LVA | 166.4 | n.d. |
| | AAV | n.d | n.d |
| | ASV | n.d | n.d |
| | ASV (chromosomal) | 176.0 | 220.2 |

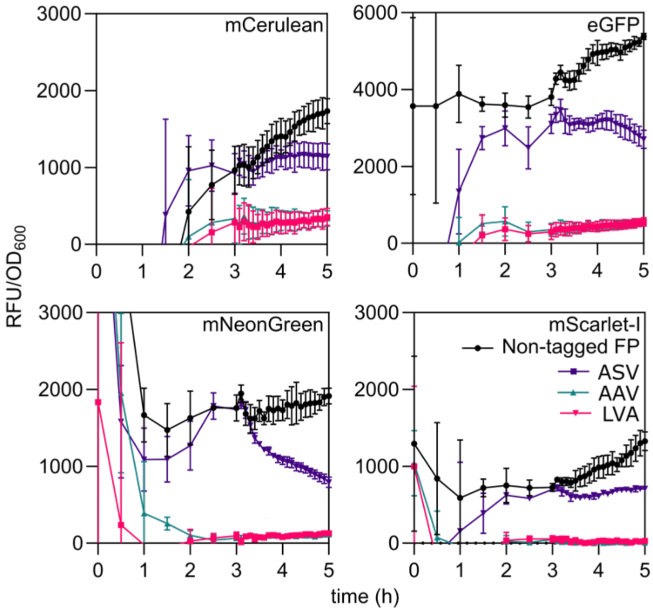


**Figure S2: Chromosomal FPs fused to SsrA degradation tags**
Strains expressing indicated FPs either non-tagged or fused to one of the three SsrA tags. Fluorescence expression was monitored over a time period of 5h and normalized to the optical density ($OD_{600}$). Depicted means and standard deviation derive of at least two biological independent experiments.

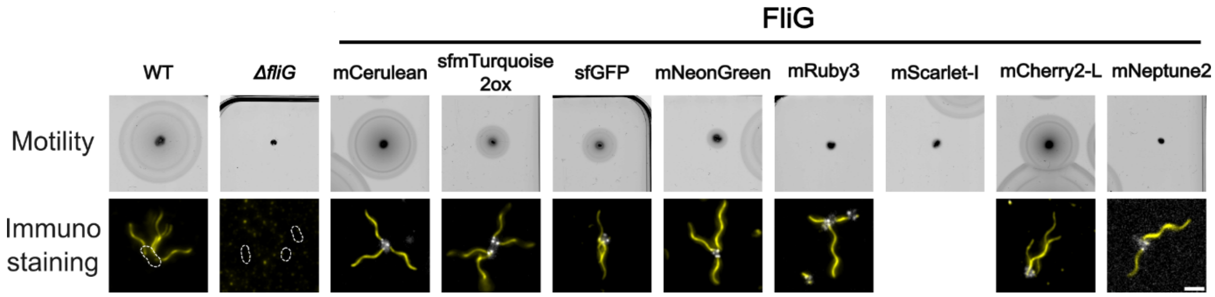


**Figure S3: Observation of swimming motility and flagellation in wild-type, Δ*fliG* and FliG N-terminal translational fusion.** Top panel: Motility assay results depicting the swimming capabilities of wild-type (WT), Δ*fliG* and FliG FPs N-terminal translational fusion strains. The motility is visualized by the expansion of the bacterial growth from a central point on a semi-solid agar plate. Bottom panel: Immunostaining images showing the flagellar filament of the strains indicated above after immunostaining of the flagellin. The images highlight the localization and distribution of the flagellar filament. Scale bar = 2 μm.

### Table S3: Strains used in this study

| Strain | Genotype | Source |
|---|---|---|
| EM7063 | LT2 / pEM2543 (pKH70-$P_{rpsM}$-*eGFP*-RPAANDENYALVA) | This study |
| EM7064 | LT2 / pEM2544 (pKH70- $P_{rpsM}$-*mNeonGreen*-RPAANDENYALVA) | This study |
| EM7065 | LT2 / pEM2545 (pKH70- $P_{rpsM}$-*mCerulean*-RPAANDENYALVA) | This study |
| EM7066 | LT2 / pEM2546 (pKH70- $P_{rpsM}$-*mScarlet-I*-RPAANDENYALVA) | This study |
| EM7067 | LT2 / pEM7067 (pKH70- $P_{rpsM}$-*eGFP*-RPAANDENYAAAV) | This study |
| EM7068 | LT2 / pEM7068 (pKH70- $P_{rpsM}$-*eGFP*-RPAANDENYAASV) | This study |
| EM7069 | LT2 / pEM7069 (pKH70- $P_{rpsM}$-*mScarlet-I*-RPAANDENYAAAV) | This study |
| EM7070 | LT2 / pEM7070 (pKH70- $P_{rpsM}$-*mNeonGreen*-RPAANDENYAASV) | This study |
| EM7071 | LT2 / pEM7071 (pKH70- $P_{rpsM}$-*mScarlet-I*-RPAANDENYAASV) | This study |
| EM7072 | LT2 / pEM7072 (pKH70- $P_{rpsM}$-*mNeonGreen*-RPAANDENYAAAV) | This study |
| EM7073 | LT2 / pEM7073 (pKH70- $P_{rpsM}$-*mCerulean*-RPAANDENYAAAV) | This study |
| EM7074 | LT2 / pEM7074 (pKH70- $P_{rpsM}$-*mCerulean*-RPAANDENYAASV) | This study |
| EM8313 | pEM8313 (pKH70-$P_{rpsM}$-*mCerulean*, AmpR) | This study |
| EM8313 | pEM8318 (pKH70-$P_{rpsM}$-*sfmTurquoise2ox*, AmpR) | This study |
| EM8314 | pEM8314 (pKH70-$P_{rpsM}$-*mRuby3*, AmpR) | This study |
| EM8315 | pEM8315 (pKH70-$P_{rpsM}$-*mCherry2-L*, AmpR) | This study |
| EM8317 | pEM8317 (pKH70-$P_{rpsM}$-*sfGFP*, AmpR) | This study |
| EM8731 | pEM8731 (pKH70-$P_{rpsM}$-*mNeonGreen*, AmpR) | This study |
| EM8733 | pEM8733 (pKH70-$P_{rpsM}$-*mCherry*, AmpR) | This study |
| EM8734 | pEM8734 (pKH70-$P_{rpsM}$-*Ypet*, AmpR) | This study |
| EM9177 | pEM9177 (pKH70-$P_{rpsM}$-*mGFPmut2*, AmpR) | This study |
| EM10686 | pEM10686 (pKH70 $P_{rpsM}$-*mScarlet-I*, AmpR) | This study |
| EM10687 | pEM10687 (pKH70-$P_{rpsM}$-*mNeptune2*, AmpR) | This study |
| EM10688 | *pEM10688 (pKH70-$P_{rpsM}$-mNeptune2.5, AmpR)* | This study |
| EM10689 | pEM10689 (pKH70-$P_{rpsM}$-*miniSOG*, AmpR) | This study |
| EM10694 | pEM10694 (pKH70-$P_{rpsM}$-*mKelly2*, AmpR) | This study |
| EM11737 | Δ*amyA::*KanSceI (ΔaaM1-Stopaa495) / pWRG730 | This study |
| EM12395 | pEM12395 (pKH70-$P_{rpsM}$-*mVenusNB*, AmpR) | This study |
| EM13792 | Δ*amyA*::$P_{rpsM}$-*eGFP* / pWRG730 | This study |
| EM13793 | Δ*amyA*::$P_{rpsM}$-*mNeonGreen* / pWRG730 | This study |
| EM13794 | Δ*amyA*::$P_{rpsM}$-*mCerulean* / pWRG730 | This study |
| EM13805 | Δ*amyA*::$P_{rpsM}$-*eGFP* | This study |
| EM13806 | Δ*amyA*::$P_{rpsM}$-*mNeonGreen* | This study |
| EM13807 | Δ*amyA*::$P_{rpsM}$-*mCerulean* | This study |
| EM13813 | Δ*amyA*::$P_{rpsM}$-*eGFP*::*KanSceI* (KanSceI replacing stop codon of *eGFP*) / pWRG730 | This study |
| EM13814 | Δ*amyA*::$P_{rpsM}$-*mNeonGreen*::KanSceI (KanSceI replacing stop codon of *mNeongreen*) / pWRG730 | This study |
| EM13815 | Δ*amyA*::$P_{rpsM}$-*mCerulean*::KanSceI (KanSceI replacing stop codon of *mCerulean*) / pWRG730 | This study |
| EM13816 | Δ*amyA*::$P_{rpsM}$-*mScarlet*-I::KanSceI (KanSceI replacing stop codon of *mScarlet*-I) / pWRG730 | This study |
| EM13825 | pEM13825 (pKH70-$P_{rpsM}$-*eGFP*) | This study |
| EM13844 | Δ*amyA*::$P_{rpsM}$-*eGFP*-RPAANDENYALVA | This study |
| EM13845 | Δ*amyA*::$P_{rpsM}$-*eGFP*-RPAANDENYAAAV | This study |
| EM13846 | Δ*amyA*::$P_{rpsM}$-*eGFP*-RPAANDENYAASV | This study |
| EM13847 | Δ*amyA*::$P_{rpsM}$-*eGFP*-AANDENYALVA | This study |
| EM13848 | Δ*amyA*::$P_{rpsM}$-*mNeonGreen*-RPAANDENYALVA | This study |
| EM13849 | Δ*amyA*::$P_{rpsM}$-*mNeonGreen*-RPAANDENYAAAV | This study |
| EM13850 | Δ*amyA*::$P_{rpsM}$-*mNeonGreen*-RPAANDENYAASV | This study |
| EM13851 | Δ*amyA*::$P_{rpsM}$-*mNeonGreen*-AANDENYALVA | This study |
| EM13852 | Δ*amyA*::$P_{rpsM}$-*mCerulean*-RPAANDENYALVA | This study |
| EM13853 | Δ*amyA*::$P_{rpsM}$-*mCerulean*-RPAANDENYAAAV | This study |
| EM13854 | Δ*amyA*::$P_{rpsM}$-*mCerulean*-RPAANDENYAASV | This study |
| EM13855 | Δ*amyA*::$P_{rpsM}$-*mCerulean*-AANDENYALVA | This study |
| EM13856 | Δ*amyA*::$P_{rpsM}$-*mScarlet*-I-RPAANDENYALVA | This study |
| EM13857 | Δ*amyA*::$P_{rpsM}$-*mScarlet*-I-RPAANDENYAAAV | This study |
| EM13858 | Δ*amyA*::$P_{rpsM}$-*mScarlet*-I-RPAANDENYAASV | This study |
| EM13859 | Δ*amyA*::$P_{rpsM}$-*mScarlet*-I-AANDENYALVA | This study |
| EM13882 | Δ*amyA*::$P_{rpsM}$-*mScarlet*-I | This study |
| EM14032 | Δ*amyA*:: $P_{rpsM}$-*mScarlet*-I / pWRG730 | This study |
| EM14033 | Δ*amyA*:: $P_{rpsM}$-*mScarlet*-I-RPAANDENYAASV / pWRG730 | This study |
| EM14958 | Δ*clpxP*::FRT / pEM13825 (pKH70-$P_{rpsM}$-*eGFP*) | This study |
| EM14959 | Δ*clpxP*::FRT / pEM2543 (pKH70-$P_{rpsM}$-*eGFP*-RPAANDENYALVA) | This study |
| EM14960 | Δ*clpxP*::FRT / pEM7067 (pKH70-$P_{rpsM}$-*eGFP*-RPAANDENYAAAV) | This study |
| EM14961 | Δ*clpxP*::FRT / pEM7068 (pKH70-$P_{rpsM}$-*eGFP*-RPAANDENYAASV) | This study |
| EM10385 | *fliG*23319 (mCerulean-FliG N-ter, SAGASA) Δ*hin*-5717::FCF (*fliC*-ON) | This study |
| EM10386 | *fliG*23267 (sfmTurquoise2ox-FliG N-ter, SAGASA) Δ*hin*-5717::FCF (*fliC*-ON) | This study |
| EM10388 | *fliG*22798 (sfGFP-fliG) Δ*hin*-5717::FCF (*fliC*-ON) | This study |
| EM10391 | *fliG*22799 (mNeonGreen-FliG) Δ*hin*-5717::FCF (*fliC*-ON) | This study |
| EM10392 | *fliG*23318 (mRuby3-FliG N-ter, SAGASA) Δ*hin*-5717::FCF (*fliC*-ON) | This study |
| EM10577 | *fliG*23343 (mScarlet-I-FliG N-ter, SAGASA) Δ*hin*-5717::FCF (*fliC*-ON) | This study |
| EM10394 | *fliG*23324 (mCherry2-L-FliG N-ter, SAGASA) Δ*hin*-5717::FCF (*fliC*-ON) | This study |
| EM10395 | *fliG*23281 (mNeptune2-FliG N-ter, SAGASA) Δ*hin*-5717::FCF (*fliC*-ON) | This study |
| EM10568 | Δ*fliG*6012::FRT Δ*hin*-5717::FCF (*fliC*-ON) | This study |
| TH5861 | Δ*hin*-5717::FCF (*fliC*-ON) | Lab collection |
| TH437 | *Salmonella enterica* serovar Typhimurium LT2 | J. Roth |

### Table S4: Oligonucleotides used in this study

| Primer | Name | Sequence (5‘-3‘) |
|---|---|---|
| 3185 | 5'-EcoRI-*mCerulean*-fw | ggacagaattcataaggaggaaaaacatATGGTTAGTAAAGGGGAG |
| 3186 | 3'-NotI-*mCerulean*-rev | ATGCCTCTAGAGCGGCC |
| 3187 | 5'-EcoRI-*mRuby3*-fw | ggacagaattcataaggaggaaaaacatATGGTGTCTAAGGGCGA |
| 3188 | 3'-NotI-*mRuby3*-rev | gtactgcggccgcTTACTTGTACAGCTCGTCCA |
| 3189 | 5'-EcoRI-*sfmTurquoise2ox* | ggacagaattcataaggaggaaaaacatatgGTGAGCAAGGGCGAGGA |
| 3190 | 3'-NotI-*sfmTurquoise2ox* | gtactgcggccgcTTATCACTTGTACAGCTCGT |
| 3191 | 5'-EcoRI-*mCherry2*-L-fw | ggacagaattcataaggaggaaaaacatATGGTCTCTAAAGGCGAG |
| 3192 | 3'-NotI-*mCherry2*-L-rev | gtactgcggccgcTTATCATTTATATAACTCGTCC |
| 3193 | 5'-EcoRI-*sfGFP*-fw | ggacagaattcataaggaggaaaaacatatgTCTAAAGGTGAAGAACTGTT |
| 3194 | 3'-NotI-*sfGFP*-rev | gtactgcggccgcTTATCATTTGTAGAGCTCAT |
| 3195 | 5'-EcoRI-*mNeonGreen*-fw | ggacagaattcataaggaggaaaaacatATGGTATCGAAGGGCGAG |
| 3196 | 3'-NotI-*mNeonGreen*-rev | gtactgcggccgcttaTCATTTATACAGTTCATCCATGCC |
| 3300 | 5'-EcoRI-*mCherry*-cloning-fw | ggacagaattcataaggaggaaaaacatATGGTTTCCAAGGGCGAGGA |
| 3301 | 3'-NotI-*mCherry*-cloning-rev | CCTTTCGTTTTATTTGATGC |
| 3302 | 5'-EcoRI-pYPet-cloning-fw | ggacagaattcataaggaggaaaaacatATGTCTAAAGGTGAAGAATT |
| 3561 | 3'-NotI-pYPet-STOP-cloning-rev | gtactgcggccgcttatcaTTTGTACAATTCATTCATAC |
| 3562 | 5'-$P^{rpsM}$-*mGFPmut2*-BamHI-fw | accgcggatccCAGATGGAGTTCTGAGGTCA |
| 3563 | 3'-*mGFPmut2*-NotI-rev | gtactgcggccgcttaTTATCATTTATATAACTCAT |
| 4058 | 5'-*mScarlet*-I-EcoRI-fw | ggacagaattcataaggaggaaaaacatATGGTGAGCAAAGGCGAAGC |
| 4059 | 3'-*mScarlet*-I-XbaI-rev | ctgagctctagattattaTTTATACAGTTCATCCATGC |
| 4059 | 3'-*mScarlet*-I-XbaI-rev | ctgagctctagattattaTTTATACAGTTCATCCATGC |
| 4060 | 5'-*miniSOG*-EcoRI-fw | ggacagaattcataaggaggaaaaacatATGGAAAAAAGCTTTGTGA |
| 4061 | 3'-*miniSOG*-NotI-rev | gtactgcggccgctcattaGCCATCCAGCTGCACGCCA |
| 4064 | 5'-*mKelly2*-EcoRI-fw | ggacagaattcataaggaggaaaaacatATGGAACTGATTAAAGAAA |
| 4065 | 3'-*mKelly2*-NotI-rev | gtactgcggccgctcattaGCGCGCCACCGCCACTTCA |
| 4066 | 5'-*mNeptune*-EcoRI-fw | ggacagaattcataaggaggaaaaacatATGGTGAGCAAAGGCGAAG |
| 4067 | 3'-*mNeptune*-NotI-rev | gtactgcggccgctcattaTTTATACAGTTCATCCATGC |
| 4797 | 5'-*mVenusNB*-EcoRI | ggacagaattcataaggaggaaaaacatATGAGCAAAGGCGAAGAACT |
| 6349 | 3‘-*amyA*-RPAANDENYAlva_rv | atttcgcttcccggcagcgctctgccgccgggaacgctcaTTATTAAGCTACTAAAGCG |
| 6350 | 5‘-*mScarlet*_RPAANDENYAlva_fw | gaaggccgccatagcaccggcggcatggatgaactgtataaaAGGCCTGCTGCAAACGAC |
| 6351 | 5‘-*mNeongreen*_RPAANDENYAlva_fw | aaagcatttaccgatgtaatgggcatggatgaactgtataaaAGGCCTGCTGCAAACGAC |
| 6352 | 5‘-*mCerulean*_RPAANDENYAlva_fw | ggcggccgggattactttggggatggacgagttatataagAGGCCTGCTGCAAACGAC |
| 6353 | 5‘-*eGFP*_RPAANDENYAlva_fw | accgccgccgggatcactctcggcatggatgaactgtataaaAGGCCTGCTGCAAACGAC |
| 6354 | 5‘-RPAANDENYAaav | AGGCCTGCAGCAAACGACGAAAACTACGCTGCAGCAGTTTAATAA |
| 6355 | 3‘-*amyA*-RPAANDENYAaav_rv | atttcgcttcccggcagcgctctgccgccgggaacgctcaTTATTAAACTGCTGCAGCG |
| 6356 | 5‘-RPAANDENYAasv | AGGCCTGCAGCAAACGACGAAAACTACGCTGCATCAGTTTAATAA |
| 6357 | 3‘-*amyA*-RPAANDENYAasv_rv | atttcgcttcccggcagcgctctgccgccgggaacgctcaTTATTAAACTGATGCAGCG |
| 6358 | 5‘-*mScarlet*_RPAANDENYAaav/asv_fw | aaggccgccatagcaccggcggcatggatgaactgtataaaAGGCCTGCAGCAAACGACG |
| 6359 | 5‘-*mNeongreen*RPAANDENYAaav/asv_fw | aagcatttaccgatgtaatgggcatggatgaactgtataaaAGGCCTGCAGCAAACGACG |
| 6360 | 5‘-*mCerulean*RPAANDENYAaav/asv_fw | ggcggccgggattactttggggatggacgagttatataagAGGCCTGCAGCAAACGACG |
| 6361 | 5‘-*eGFP*_RPAANDENYAaav/asv_fw | ccgccgccgggatcactctcggcatggatgaactgtataaaAGGCCTGCAGCAAACGACG |
| 6362 | 3‘-*amyA*-AANDENYAlva_rv | atttcgcttcccggcagcgctctgccgccgggaacgctcaTTATCACGCCACCAGCGCAT |
| 6363 | 5‘-*mScarlet*_AANDENYALVA_fw | gaaggccgccatagcaccggcggcatggatgaactgtataaaGCGGCGAACGATGAAAAC |
| 6364 | 5‘-*eGFP*_AANDENYALVA_fw | accgccgccgggatcactctcggcatggatgaactgtataaaGCGGCGAACGATGAAAAC |
| 6365 | 5‘-*mNeongreen*_AANDENYALVA_fw | aaagcatttaccgatgtaatgggcatggatgaactgtataaaGCGGCGAACGATGAAAAC |
| 6366 | 5‘-*mCerulean*_AANDENYALVA_fw | ggcggccgggattactttggggatggacgagttatataagGCGGCGAACGATGAAAAC |
| 6414 | 3'-XbaI-RPAANDENYAlva_rv | tttgatgcctctagaTTATTAAGCTACTAAAGCG |
| 6415 | 3'-XbaI-RPAANDENYAaav_rv | tttgatgcctctagaTTATTAAACTGCTGCAGCG |
| 6416 | 3'-XbaI-RPAANDENYAasv_rv | tttgatgcctctagaTTATTAAACTGATGCAGCG |
| 6417 | 3'-XbaI-AANDENYAlva_rv | tttgatgcctctagaTTATCACGCCACCAGCGC |
